# A yeast model of the ALS protein Matrin3 uncovers Hsp90 and its co-chaperone Sti1 as modifiers of misfolding and toxicity

**DOI:** 10.1101/2021.08.24.457481

**Authors:** Sonja E. Di Gregorio, Mohammad Esmaeili, Ahmed Salem, Martin L. Duennwald

## Abstract

The MATR3 gene encoding the protein Matrin3 is implicated in the pathogenesis of the neurodegenerative disease amyotrophic lateral sclerosis (ALS). Matrin3 forms neuronal cytoplasmic and nuclear inclusions in ALS-affected neurons. Additionally, 13 heterozygous missense mutations in MATR3 are identified in ALS patients. To further explore Matrin3 misfolding and toxicity, we established and characterized a yeast model. We demonstrate that wild type Matrin3 and the ALS-associated variant F115C are toxic and form inclusions in yeast. Our further characterization uncovers substantial modification of Matrin3 toxicity and inclusion formation by Hsp90 and its co-chaperones, specifically Sti1. Thus, our study demonstrates how specific branches of cellular protein quality control regulate the misfolding and toxicity of Matrin3.

**Summary Statement:** We established and characterized a yeast model expressing human Matrin3, a protein implicated in the pathogenesis of amyotrophic lateral sclerosis (ALS). Using this yeast model and mammalian neuronal cells, we showed that Matrin3 mislocalizes and forms inclusions, is cytotoxic, and increases sensitivity to cellular stress. We also uncovered that Hsp90 and particularly its co-chaperone Sti1 alter Matrin3 toxicity.

## Introduction

Degeneration of the upper and lower motor neurons of the spinal cord and brain characterize amyotrophic lateral sclerosis (ALS) (Hardiman et al., 2017), leading to a progressive loss of muscle function and paralysis resulting in death often 1-2 years following diagnosis (Polkey et al., 1999, Leigh et al., 2003). The majority of ALS cases are characterized as sporadic (∼90%, sALS), where the underlying causes are unclear. ALS cases with a pre-existing family history are termed familial (∼5-10% of cases, fALS) (Chen et al., 2013). Mutations in more than 18 different genes are currently known to cause fALS, with gene discovery rapidly increasing the number of affected genes since the first ALS-associated mutations in SOD1 were identified (Srinivasan and Rajasekaran, 2020, Gitler and Fryer, 2018). Many of the proteins encoded by known ALS genes are involved in RNA metabolism and share similarities in ALS pathogenesis. Protein misfolding and mislocalization are common features of ALS, with many ALS proteins found mislocalized from functional compartments and sequestered into inclusions in the cytosol and nucleus (Parakh and Atkin, 2016, Blokhuis et al., 2013). Despite considerable research efforts, the molecular mechanisms underpinning ALS remain mostly unknown, and there are no effective treatments.

Protein misfolding is a global hallmark of neurodegenerative disorders (Parakh and Atkin, 2016, Soto and Estrada, 2008, Soto, 2003). Changes in the structure of a soluble protein that lead to compromised function, structure, and often localization are considered protein misfolding. (Hartl, 2017). Misfolded ALS proteins, such as TDP-43, FUS, and SOD1, mislocalize and form pathological inclusions, mostly in the cytoplasm of motor neurons. These are termed neuronal cytoplasmic inclusions (NCIs) and are a pathological hallmark of ALS (Saberi et al., 2015). Genetic mutations can increase the propensity of ALS proteins to misfold. However, even wild type variants of ALS proteins, such as TDP-43, are found mislocalized to NCIs in sALS (Kryndushkin and Shewmaker, 2011, Redler and Dokholyan, 2012, Patel et al., 2015). Environmental insults, such as changes in pH, exposure to toxic chemicals, excessive oxidative stress, or aging can also contribute to protein misfolding and neurodegeneration in ALS (Cannon and Greenamyre, 2011, Barber et al., 2006, Kikis et al., 2010).

Matrin3 is a 130kDa RNA and DNA binding protein encoded by the *Matr3* gene in humans (Belgrader et al., 1991), which is highly conserved in metazoans (Belgrader et al., 1991, Nakayasu and Berezney, 1991, Hibino et al., 2006). Matrin3 is a nuclear matrix protein containing two characteristic C_2_H_2_ Zink-finger binding motifs (ZnF1 and ZnF2), also referred to as ‘Matrin homology domains,’ two tandem RNA recognition motifs (RRM1 and RRM2), and a nuclear localization signal (NLS) and nuclear export signal (NES) (Nakayasu and Berezney, 1991, Belgrader et al., 1991, Hibino et al., 2006, Hisada-Ishii et al., 2007). Matrin3 is ubiquitously expressed and found at high levels in neurons compared to other cell types in the human central nervous system (Verkerk et al., 1991). It is localized throughout the nucleoplasm and enriched at the inner nuclear matrix. Along with other proteins, Matrin3 forms a nuclear scaffolding network required for genetic regulatory processes and chromatin organization (Malyavantham et al., 2008, Zeitz et al., 2009). In addition, Matrin3 regulates RNA processing, DNA replication, transcription, and is implicated in repairing DNA double-strand breaks and NMDA-induced neuronal death (Salton et al., 2010, Giordano et al., 2005, Hibino et al., 2006, Hibino et al., 1992). Matrin3 participates in alternative splicing events mostly independent of the splicing regulator PTB (Coelho et al., 2015). Accordingly, Matrin3 interacts with proteins that function in RNA processing (Gallego-Iradi et al., 2015, Coelho et al., 2015). Many of the known interactors of Matrin3 are RNA binding proteins, and Matrin3 stabilizes 77 different mRNA transcripts (Salton et al., 2011). Matrin3 has also been implicated in the retention of hyper-edited viral mRNA in the nucleus, thereby preventing their translation (Zhang and Carmichael, 2001).

Matrin3 is essential for the viability of specific cell types and mutations in Matrin3 have been linked to asymmetric distal myopathy with vocal cord paralysis and ALS (Hisada-Ishii et al., 2007, Przygodzka et al., 2011, Johnson et al., 2014, Muller et al., 2014, Senderek et al., 2009) Further, alterations in Matrin3 levels are implicated in fetal Down Syndrome and macronodular adrenocortical disease and adrenocortical tumorigenesis (Bimpaki et al., 2010, Bernert et al., 2002). A total of 13 mutations have been identified in Matrin3 that cause ALS with the S85C mutant also found in distal myopathy (Johnson et al., 2014, Boehringer et al., 2017, Senderek et al., 2009, Lin et al., 2015, Xu et al., 2016, Marangi et al., 2017, Leblond et al., 2016, Origone et al., 2015). ALS-associated amino acid substitutions in Matrin3 are found in the protein regions that are predicted to be intrinsically disordered. (Malik et al., 2018). Pathologically, studies in neurons of ALS patients’ spinal cord and brain tissues demonstrate that mutant variants and wild type Matrin3 mislocalizes to NCIs and show abnormally strong staining within the nucleus (Tada et al., 2018, Johnson et al., 2014). Mislocalization and strong staining in the nucleus was also observed for Matrin3 in patients with *C9orf72* hexanucleotide expansions and *FUS* mutations (Dreser et al., Tada et al., 2018, Johnson et al., 2014). Thus, Matrin3 joins a growing list of ALS proteins involved in RNA processing (TDP-43, FUS, C9ORF72, ATXN2, TAF15, hnRNPA1, hnRNPB1, EWSR1, SETX). Yet, these underlying mechanisms by which Matrin3 contributes to neurodegeneration in ALS remain mostly unclear (La Cognata et al., 2020).

Cellular protein quality control and molecular chaperones modulate protein misfolding in neurodegenerative disorders, including ALS (Meriin and Sherman, 2005, Ciechanover and Kwon, 2017, Sweeney et al., 2017). Heat shock proteins (Hsps) and molecular chaperones facilitate the unfolding and refolding of proteins and prevent misfolding and aggregation. Matrin3 has been previously reported to interact with the heat shock proteins glucose-regulated protein 78 (GRP78), GRP75, and glutathione S-transferase π isoform 2 (GSTπ2), which regulate Matrin3 degradation (Osman and van Loveren, 2014). Yet, how other molecular chaperones modulate Matrin3 misfolding and toxicity remains unknown. Importantly, the molecular chaperone networks are highly conserved between humans and yeast, making yeast an excellent model organism to study these processes in a genetically and biochemically tractable model.

Yeast models expressing misfolded proteins associated with neurodegenerative diseases, including ALS, helped decipher key proteins and mechanisms (Di Gregorio and Duennwald, 2018b, Di Gregorio and Duennwald, 2018a, Krobitsch and Lindquist, 2000, Hofer et al., 2018) that contribute to neurodegeneration. Experiments in yeast are highly informative for exploring the interaction between the highly conserved cellular protein quality control and protein misfolding (Fields and Johnston, 2005). Here, we established and characterized a yeast model expressing human wild type Matrin3 and the ALS variant F115C and employed cultured neuronal cells to explore Matrin3 misfolding, toxicity, and its interactions with specific branches of cellular protein quality control. In parallel to many other yeast models, Yeast cells do not express any close homologue of Matrin3, thus allowing focused studies on its toxicity independent of its regular function. Our results from both yeast and mammalian neuronal cells document that Hsp90 and its co-chaperones Sti1, alter Matrin3 misfolding and toxicity.

## Results

First, we established and characterized a yeast model expressing human Matrin3. Figure 1A shows a schematic representations of the domain structure of wild type Matrin3 and all variants used in our study. We found that high expression (Gal promotor, 2μ i.e. multi-copy plasmid) of wild type Matrin3 almost completely blocks yeast growth (Figure 1B, non-inducing control plate available in Supplementary figure 7A). High expression of the ALS variant F115C is as toxic as wild type Matrin3 (Figure 1B). We monitored the subcellular localization of Matrin3 in yeast by expressing yellow fluorescent protein (YFP) carboxy-terminal fusion to full length Matrin3 (Matrin3-YFP) and F115C variant (F115C-YFP, Figure 1C). Wild type Matrin3-YFP and F115C-YFP form fluorescent foci in yeast cells (Figure 1C, expanded fields available in Supplementary figure 1B). Wild type Matrin3-YFP and F115C-YFP are found exclusively localized in foci, with some cells displaying multiple foci.

**Figure 1.**
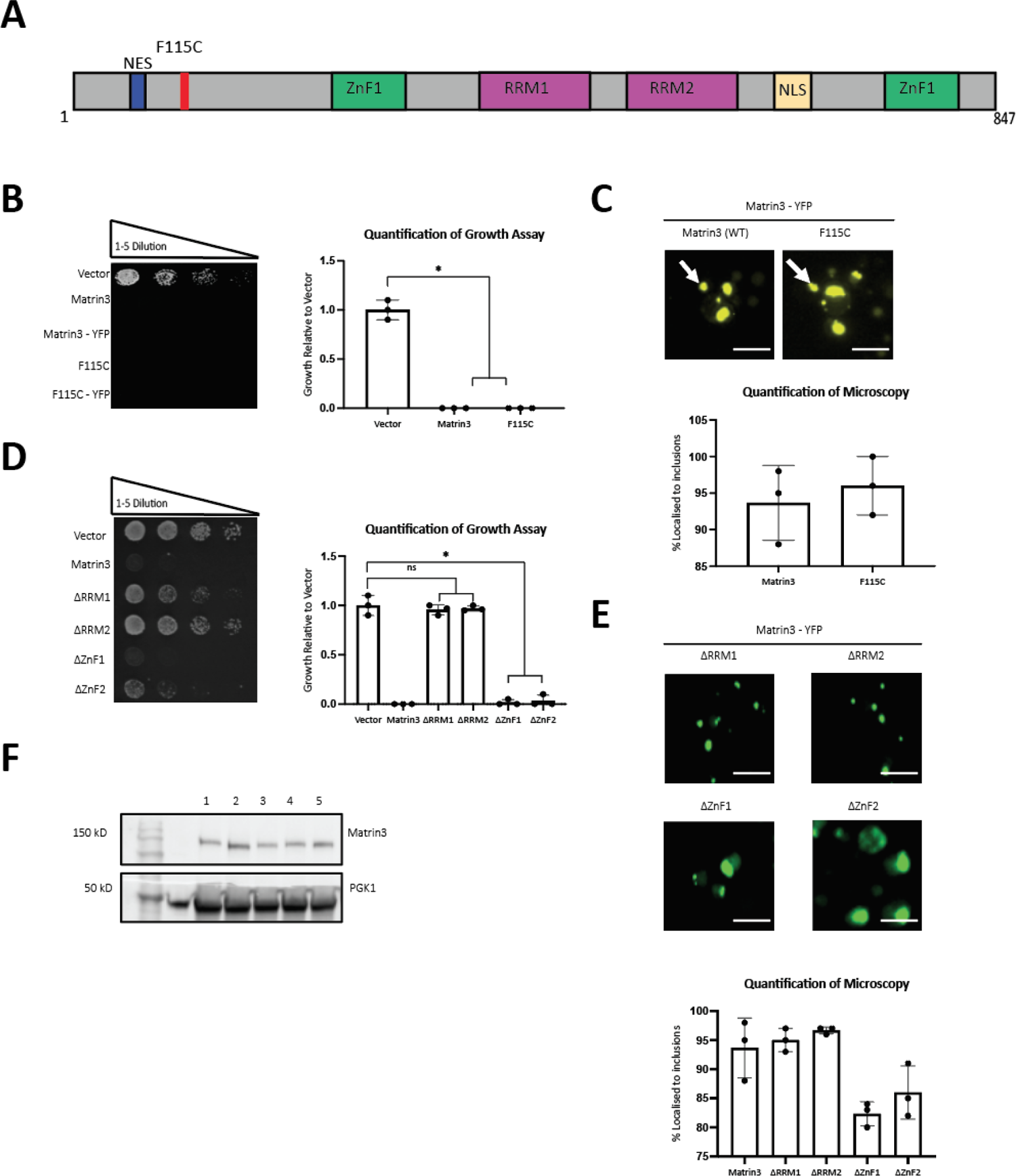
The Matrin3 yeast model. (**A**) Schematic representation of full length wild type Matrin3, its functional domains, and the F115C amino acid substitution. (**B**) Growth assay of yeast cells expressing human wild type Matrin3 and F115C (high expression system). Both untagged and - Yellow Fluorescent Protein (YFP) tagged constructs were evaluated. Results were quantified and normalized to growth of the vector control harboring cells. Error bars represent standard deviations. (* = *p* < 0.01). **(C)** Matrin3-YFP and F115C–YFP localization and inclusion formation was observed by fluorescence microscopy in live yeast cells. Arrows indicate inclusions. The scale bar corresponds to 5µm. Bar graph represents the quantification of data as a percentile of cells containing inclusions. **(D)** Growth assay of yeast cells expressing the indicated Matrin3 domain truncated variants. (* = *p* <0.01). **(E)** Matrin3 truncated variants-YFP localization and inclusion formation was observed by fluorescence microscopy in live yeast cells. The scale bar corresponds to 5µm. Bar graph represents the quantification as a percentile of cells containing inclusions. **(F)** Expression of Matrin3 constructs (untagged) are confirmed by Western blot probed with an anti-Matrin3 antibody. PGK1 serves as loading control.

We next expressed domain-truncated variants of Matrin3, each lacking one of the tandem RNA recognition motifs (RRM1 or RRM2) or one of the Zinc-Finger binding motifs (ZnF1 or ZnF2) (Malik et al., 2018) in yeast. Toxicity equivalent to WT Matrin3 is observed when the truncation lacking either ZnF domains are expressed in yeast (Figure 1D, non-inducing control plate available in Supplementary figure 7A). However, truncations of either of the two RRM domains almost completely rescues growth (Figure 1D). Our data in yeast thus confirm results in mammalian cells and *Drosophila*, indicating that the RRM domains contribute to Matrin3 toxicity (Ramesh et al., 2020, Malik et al., 2018). We found that all truncation variants fused to YFP showed inclusions in most cells (Figure 1E, expanded fields available in Supplementary figure 1C). Western blot analysis of wild type Matrin3 and truncated variants confirm comparable steady-state protein levels (Figure 1F). To determine if the foci observed are biochemically insoluble, such as polyglutamine proteins or some yeast prions (Halfmann and Lindquist, 2008, Scherzinger et al., 1997), or soluble, such as many ALS proteins, including TDP-43 and FUS (Johnson et al., 2008, Shelkovnikova et al., 2014), we performed semi denaturing detergent gel electrophoresis (SDD-AGE) with lysates prepared for yeast expressing Matrin3 and Matrin3-YFP. Lysates from yeast cell expressing huntingtin fragments served as controls (Supplementary Figure 1A). 25Q is non-toxic in yeast and does not form aggregates and presents only a single band (soluble monomer) on the SDD-AGE assay, whereas 72Q is toxic and forms insoluble aggregates, which runs as a high-molecular weight smear (insoluble oligomers) and single band (soluble monomer) as reported before (Scherzinger et al., 1997, Halfmann and Lindquist, 2008). SDD-AGE of Matrin3 and Matrin3-YFP thus shows that the microscopically detectable Matrin3-YFP inclusions are soluble, similarly to TDP-43 and FUS (Guo et al., 2011, Bosque et al., 2013). We also observed some possible proteolysis of Matrin3 indicated by faster migrating protein species in our SDD-AGE analysis (Supplementary Figure 1A).

We also tested whether exposure to cellular stress modulates Matrin3 misfolding and toxicity in yeast. We used AZC (a proline analogue), MG132 (a proteasome inhibitor), or radicicol (an Hsp90 inhibitor) to induce protein misfolding and quality control stress, tunicamycin or DTT to induce endoplasmic reticulum (ER) stress, and hydrogen peroxide to induce oxidative stress (Trotter et al., 2001, Ling and Söll, 2010, Dukan et al., 2000, Braakman et al., 1992, Schulte et al., 1999). We performed these growth assays with a low constitutive expression system (GPD promotor, CEN plasmid, i.e. single plasmid per cell) for Matrin3 to allow detection of increased toxicity. Results were quantified and normalized to growth of the control cells. Lower expression of Matrin3 reduced toxicity by ∼50% in yeast compared high expression (Figure 2A, non-inducing control plate available in Supplementary figure 7B). Western blot analysis of Matrin3 expression under the constitutive (GPD) promotor show ∼75% lower levels of protein in cells (Figure 2I).

**Figure 2.**
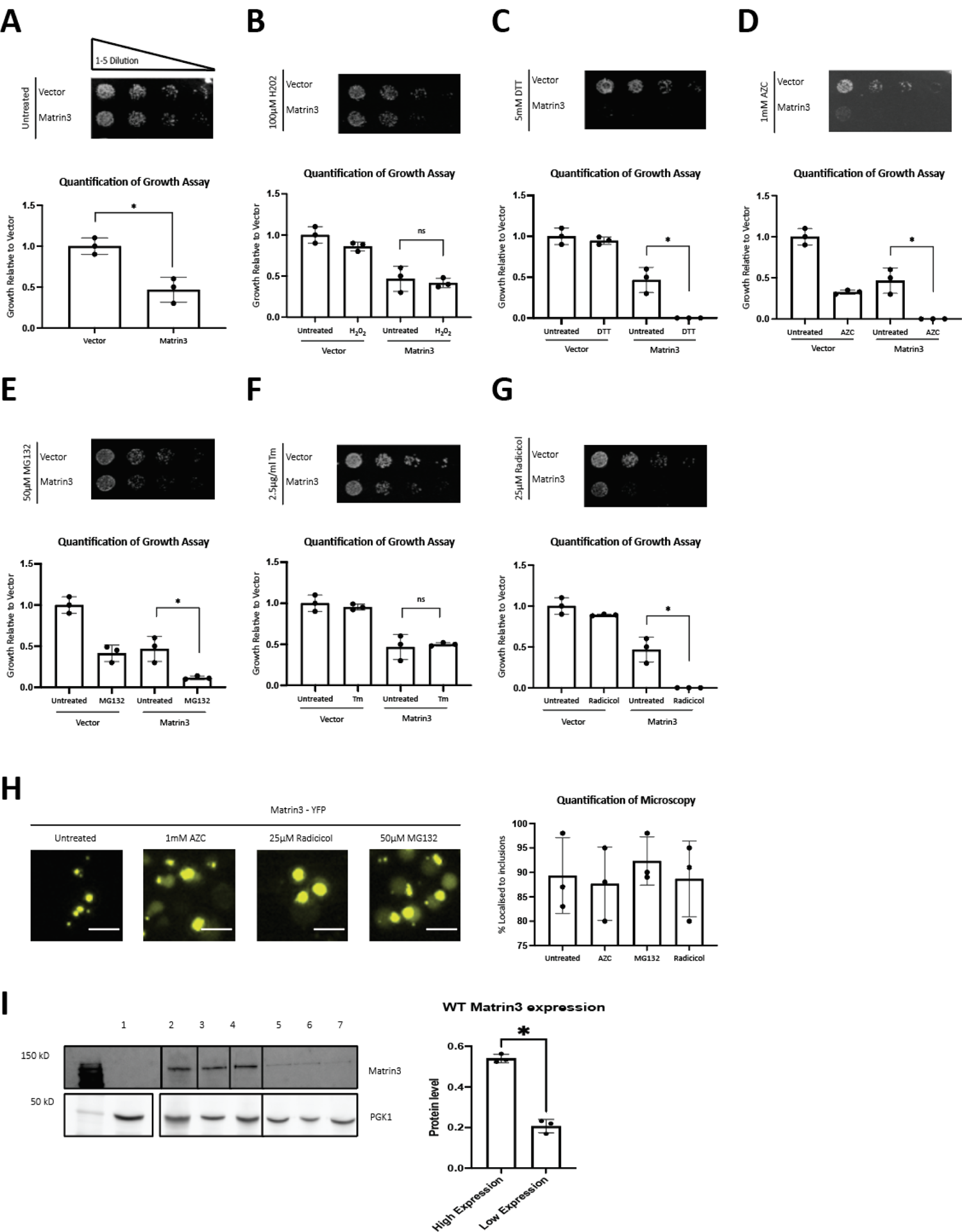
Protein folding stress exacerbates Matrin3 toxicity. (**A**) Growth assay of yeast cells expressing Matrin3 (low expression system) was evaluated. Results were quantified and normalized to growth of the control cells transformed with empty vector, (* = *p* < 0.05). Error bars represent standard deviations. Growth assays of yeast cell expressing Matrin3 (low expression system) on media containing **(B)** 100µM H_2_0_2_, **(C)** 5mM DTT, (**D)**1mM AZC, **(E)** 50µM MG132 (here a yeast strain deleted for PDR5 was used to enable MG132 activity, **(F)** 2.5µg/ml tunicamycin (Tm), and **(G)** 25µM radicicol. **(H)** Matrin3-YFP localization and inclusion formation in the presence of the indicated stress conditions was observed using fluorescence microscopy in live yeast cells. The scale bar corresponds to 5µm. Bar graph represents the quantification as percentile of cells containing inclusions. All error bars represent standard deviations (* = *p* <0.05). (I) Western blot analysis of Matrin3 expressed at high and low levels.

Compared to Matrin3 expressing cells grown on control plates, cells expressing Matrin3 grown on media containing hydrogen peroxide did not show altered toxicity (Figure 2B). Cells expressing Matrin3 grown in the presence of the ER stressor DTT but not tunicamycin show enhanced toxicity (Figure 2C, 2F). Matrin3 toxicity was exacerbated by ACZ, MG132, and radicicol (Figure 2D-E, G). Finally, Figure 2H shows that Matrin3-YFP localization and inclusion formation appears unchanged in the presence of all stressors tested here (expanded fields available in Supplementary figure 1C).

Following our studies in Hsp90, its co-chaperones ad TDP-43, (Lin et al., 2021), we explored if Hsp90 and its co-chaperones modulate Matrin3 toxicity and inclusion formation. Yeast express two Hsp90 paralogues, Hsc82 and Hsp82 (Mortimer et al., 1992, Nathan et al., 1997, Mortimer et al., 1989). Deletion of Hsc82 greatly enhances Matrin3 toxicity (GPD promotor, CEN plasmids) compared to WT (Figure 3A, non-inducing control plate available in Supplementary figure 7C). In addition, deletion of HSC82 greatly reduced Matrin3-YFP inclusion formation (Figure 3C, expanded fields available in Supplementary figure 3A). Similarly, deletion of HSP82 also enhances Matrin3 toxicity and decreases Matrin3-YFP inclusion formation, albeit to a lesser extent than deletion of HSC82 (Figure 3A, C, non-inducing control plate available in Supplementary figure 7C). This data document that reducing Hsp90 levels alter both Matrin3 inclusion formation and toxicity in yeast.

**Figure 3.**
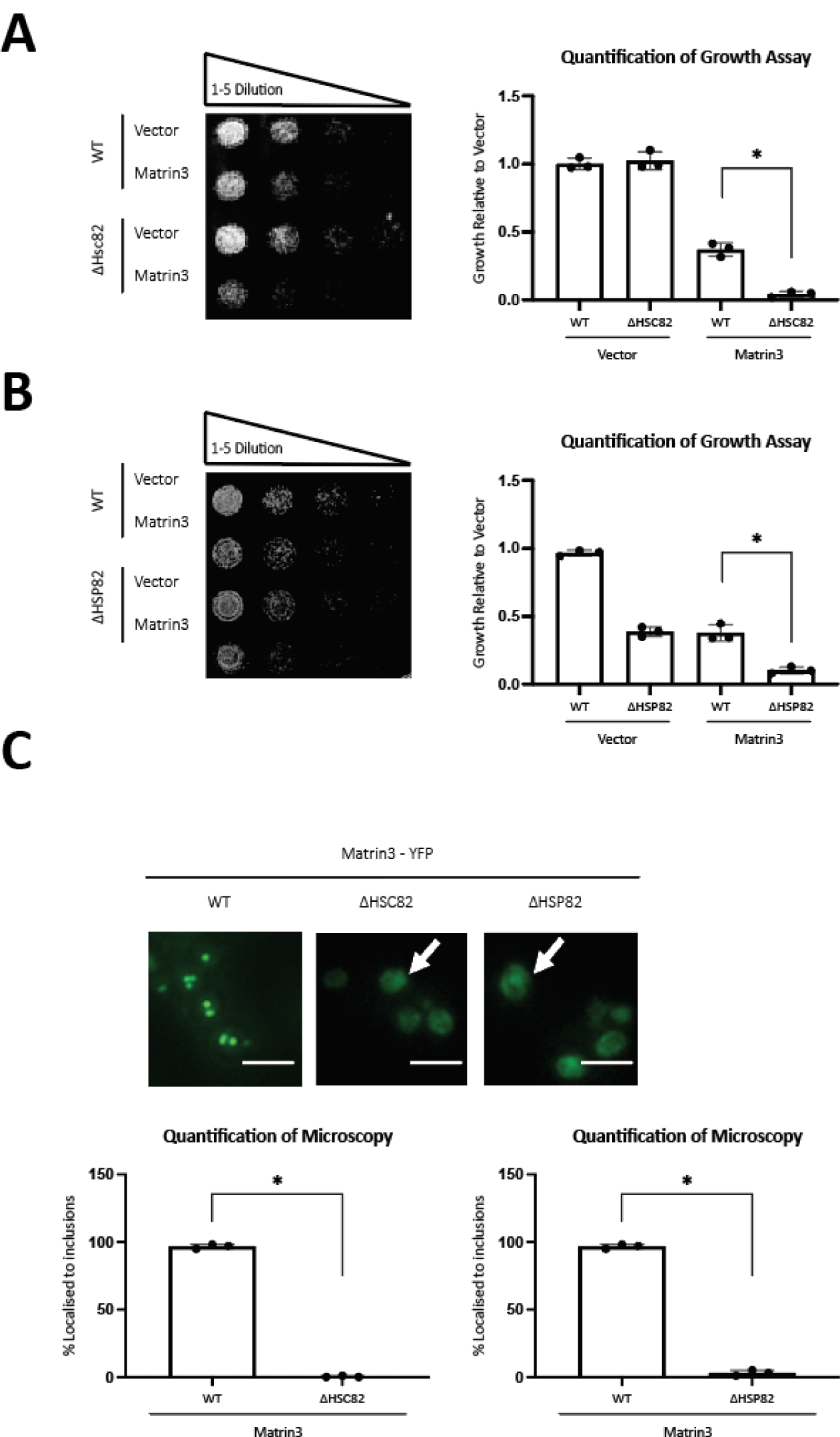
Matrin3 toxicity and inclusion formation are altered by HSP90 deletion. Growth assay of yeast cells expressing Matrin3 (low expression system) in wild type and **(A)** Δhsc82 and **(B)** Δhsp82 strains. Results were quantified and normalized to growth of the wild type strain vector control cells. Error bars represent standard deviations. (* = *p* < 0.05). **(C)** Matrin3-YFP localization and inclusion formation was observed using fluorescence microscopy in wild type and Hsp90 deletion strains. The scale bar corresponds to 5µm. Bar graph represents the quantification of data as a percentile of cell populations containing inclusions (* = *p* < 0.01).

We next asked if deletion of Hsp-90 co-chaperones affects Matrin3 cytotoxicity and inclusion formation. We first tested the co-chaperones Sti1 and Aha1. Sti1 (Hop in mammals), is a stress inducible co-chaperone that facilitates client protein transfer between Hsp70 and Hsp90 (Nicolet and Craig, 1989, Chang et al., 1997, Richter et al., 2003, Li et al., 2011, Wolfe et al., 2013). Aha1 binds to and stimulates intrinsic ATPase activity of Hsp90 leading to conformational changes in the catalytic loop (Panaretou et al., 2002, Lotz et al., 2003). When Matrin3 is expressed in yeast cells deleted for STI1, its toxicity is enhanced compared to wild types cells (Figure 4A, non-inducing control plate available in Supplementary figure 7D). The deletion of AHA1 also exacerbated Matrin3 toxicity (Figure 4B, non-inducing control plate available in Supplementary figure 7D). When either co-chaperone is absent, Matrin3-YFP is mostly diffusely localized throughout the cytoplasm and inclusion formation is reduced by ∼75% compared to wild type cells (Figure 4C, Supplementary Figure 3B). We next examined Matrin3 growth in yeast strains with reduced expression levels (DAmP, Decreased Abundance by mRNA Perturbation) of essential Hsp90 co-chaperones, Cdc37, Ess1, and Sgt1. Cdc37 interacts with Hsp82 directly and aids in the protein refolding process (Abbas-Terki et al., 2002, Mandal et al., 2007). Ess1 is a Peptidylprolyl-cis/trans-isomerase (PPIase) specific for phosphorylated S/T residues N-terminal to proline (Hanes et al., 1989). Sgt1 is upregulated during DNA replication stress and is a co-chaperone that links Skp1p to Hsp90 complexes (Catlett and Kaplan, 2006, Tkach et al., 2012). Reduced expression of all three Hsp90 co-chaperones enhanced Matrin3 toxicity compared to the wild type strain (Supplementary Figure 3A, non-inducing control plate available in Supplementary figure 7E). DAmP alleles of all these Hsp90 co-chaperones greatly reduced Matrin3 inclusion formation in yeast (Supplementary Figure 2A, B).

**Figure 4.**
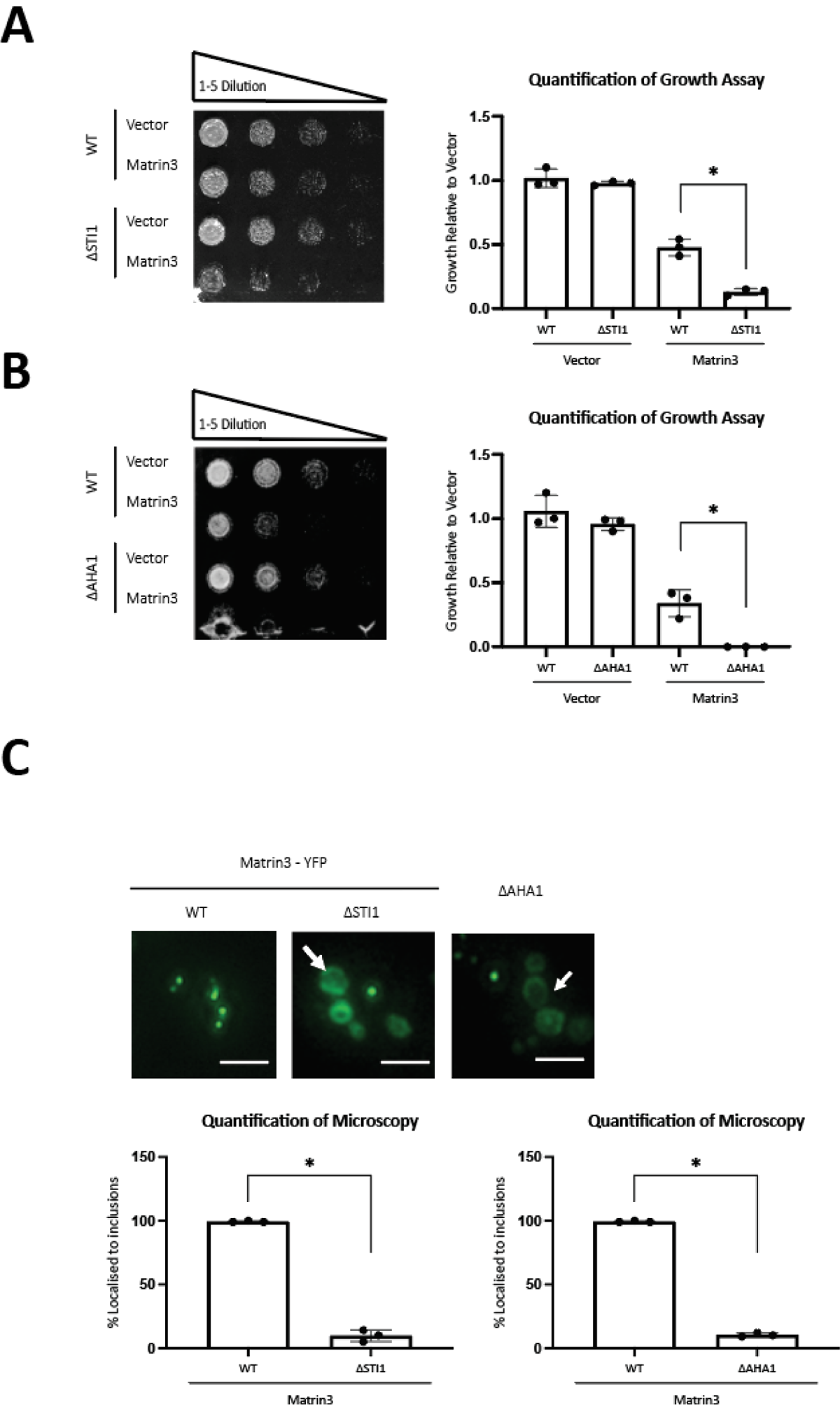
Matrin3 toxicity and inclusion formation are altered by Hsp90 co-chaperone deletions. **(A)** Growth assay of Matrin3 expressing (low expression system) wild type and Δsti1 cells. Spotting assay were quantified and normalized to growth of the wild type strain vector control cells. (* = *p* < 0.05). (**B**) Growth assay of yeast cells expressing Matrin3 (low expression system) in wild type and Δaha1 cells and quantification. (* = *p* < 0.01). **(C)** Matrin3-YFP localization and inclusion formation was observed by fluorescence microscopy in wild type and Δhsc82, Δhsp82, and Δaha1 cells. Arrows indicate diffuse Matrin3 localization. The scale bar corresponds to 5µm. Bar graph represents the quantification of cells containing inclusions. (* = *p* < 0.05).

We examined how overexpression of Hsp90 and its co-chaperones modulates Matrin3 toxicity and inclusion formation. Overexpression of Aha1 and Hsc82 enhanced Matrin3 toxicity (Figure 5C, non-inducing control plate available in Supplementary figure 7F) but inclusion formation was not altered (Supplementary Figure 3D). Moderate Sti1 overexpression (GPD promotor, CEN plasmid) reduces Matrin3 toxicity (Figure 5A, non-inducing control plate available in Supplementary figure 7F), whereas high Sti1 overexpression (GAL promotor, 2micron plasmid) is toxic to cells generally and enhances Matrin3 toxicity (Figure 5B, non-inducing control plate available in Supplementary figure 7F). Sti1 overexpression does not affect Matrin3 inclusion formation (Supplementary Figure 3C).

**Figure 5.**
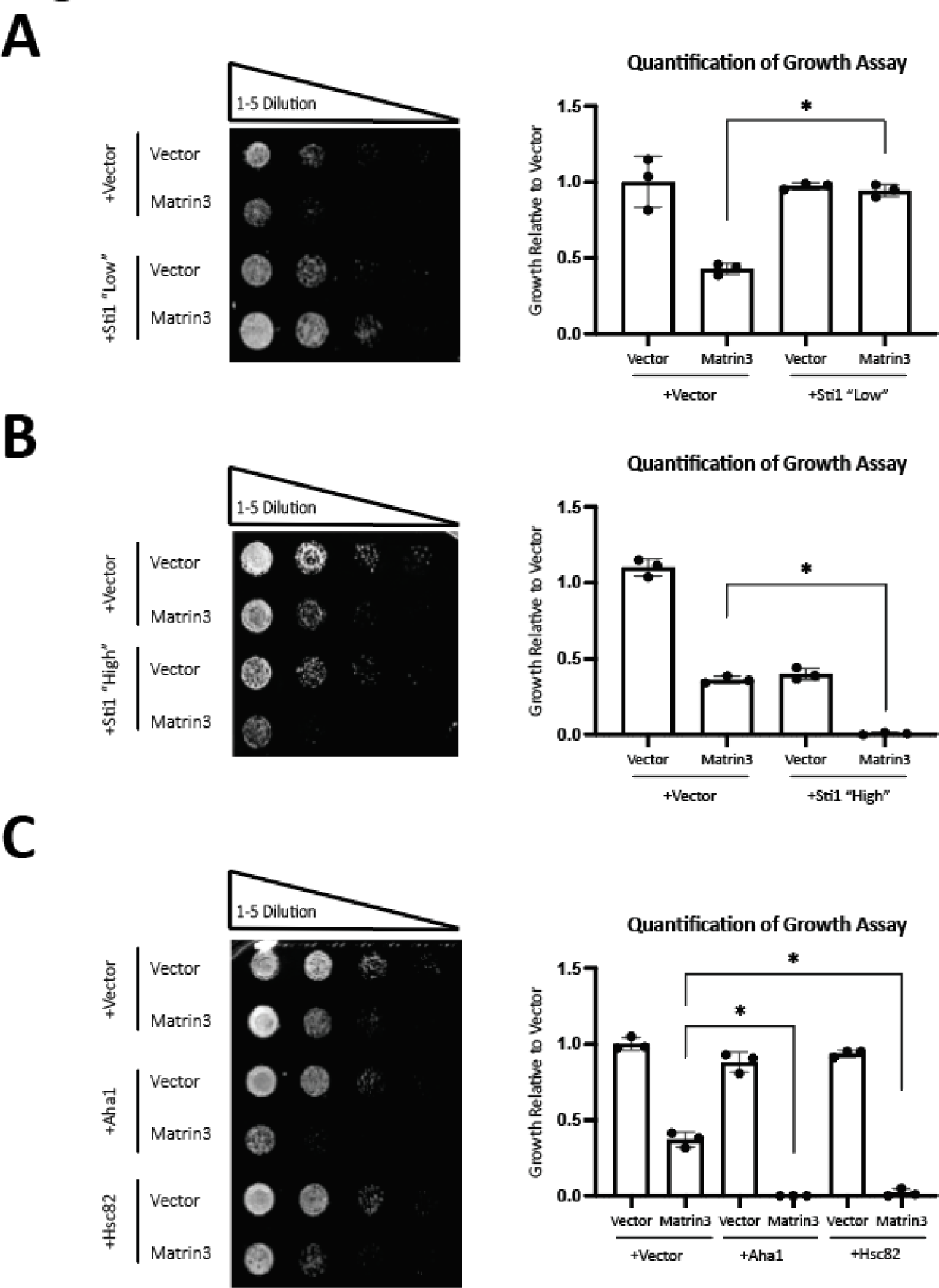
Matrin3 toxicity and inclusion formation are altered by overexpression of Hsp90 and Hsp90 co-chaperones. **(A)** Growth assay of Matrin3 expressing yeast cells (low expression system) overexpressing Sti1 (* = *p* < 0.05). **(B)** Matrin3 yeast growth assay (low expression) in wild type and overexpressed (high expression, i.e. 2μ plasmid) Sti1 strains. Results were quantified and normalized to growth of the wild type strain vector control cells (* = *p* < 0.05). **(C)** Matrin3 yeast growth assay (low Matrin3 expression system) in WT and overexpressed Aha1 or Hsc82 strains. * = *p* < 0.05).

To validate our findings obtained using the Matrin3 yeast model in mammalian neuronal cells we utilized two murine cell lines, cholinergic SN56 and neuroblastoma-derived Neuro-2a cells (N2a cells, Supplementary Figure 4) both of which are well-established models to study protein misfolding (Hammond et al., 1990, LePage et al., 2005). First, we assessed the toxicity associated with Matrin3 overexpression in SN56 cells transiently transfected with Matrin3 expression vectors. Figure 6A shows overexpression of Matrin3 in SN56 cells is mildly toxic as assessed by a viability assays that measure ATP levels. This toxicity is exacerbated by oxidative stress (hydrogen peroxide) and protein misfolding stress (MG132). Inhibition of Hsp90 by radicicol does not increase the toxicity associated with overexpressed Matrin3 (Figure 6A). Further, immunofluorescence confocal microscopy of Matrin3 in SN56 and N2a (Supplementary Figure 4 and 5) cells treated with radicicol and MG132 show an increase in Matrin3 inclusions in the cytoplasm compared to untreated cells (Figure 6B), whereas hydrogen peroxide treatment had no effect on inclusions. Three distinct localization patterns were observed and quantified in Figure 6C; (1) Foci, (2) Nuclear overabundance with cytosolic foci and (3) diffuse. Untreated cells display diffuse (pattern 3) Matrin3 localization in the nucleus. Treatment with MG132 leads to the formation of foci in the majority of cells. Radicicol leads to an increase in the proportion of cells displaying pattern 1 and 2. Overexpression of Matrin3 leads to a higher proportion of cells showing diffuse localization in the nucleus compared to endogenous expression of Matrin3 when treated with radicicol. Figure 6D shows higher levels of overexpressed Matrin3 compared to endogenous Matrin3 (Figure 6D).

**Figure 6.**
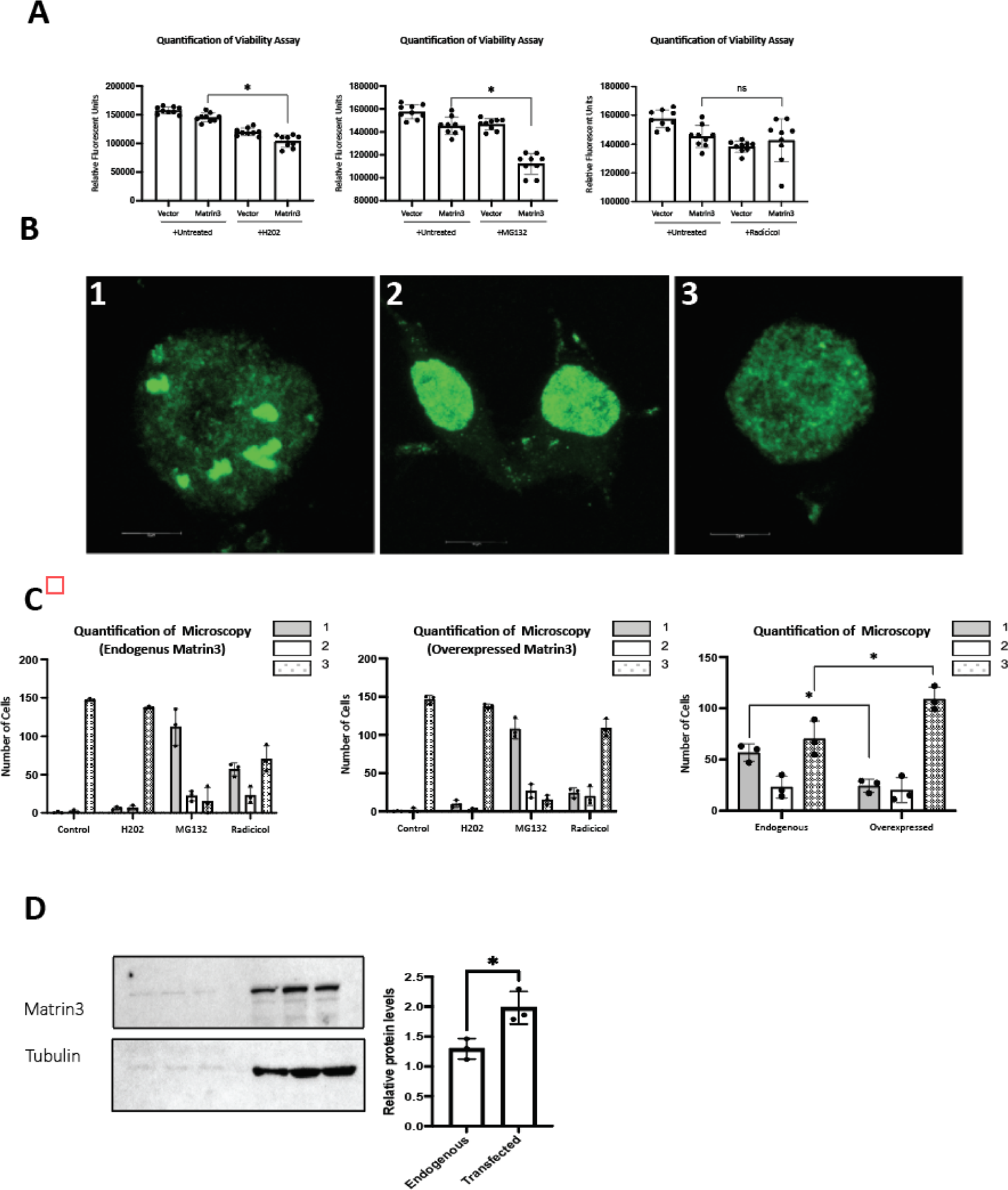
Matrin3 toxicity in mammalian cells is exacerbated by protein folding stress. **(A)** ATP-viability assays SN56 cells transiently transfected with expression vectors for WT Matrin3 and vector controls (* = *p* < 0.05) a treated with MG132 or H_2_0_2_ (* = *p* >.01), and radicicol. **(B)** Immunofluorescence microscopy of endogenous and overexpressed (transiently transfected) Matrin3 in SN56 cells in untreated and cells treated with MG132, H_2_0_2_, and radicicol. Scale bars correspond to 100µM. **(D)** Western blot analysis of endogenous and overexpressed Matrin3 in SN56 cells.

We also explored the effect Sti1 on Matrin3 toxicity and inclusion formation in mammalian neuronal cells. Overexpression of Sti1 simultaneously with radicicol treatment rescues Matrin3 toxicity (Figure 7A) compared to radicicol treatment alone. To assess Matrin3 toxicity in cells deleted for Sti1, we transiently transfected SN56 Sti1 knockout cells with Matrin3 expression vectors (Lackie et al., 2020). Matrin3 toxicity is enhanced by Hsp90 inhibition (radicicol) in the Sti1 knock out cells when compared to Matrin3 toxicity in the Sti1 knockout line without treatment (Figure 7B).

**Figure 7.**
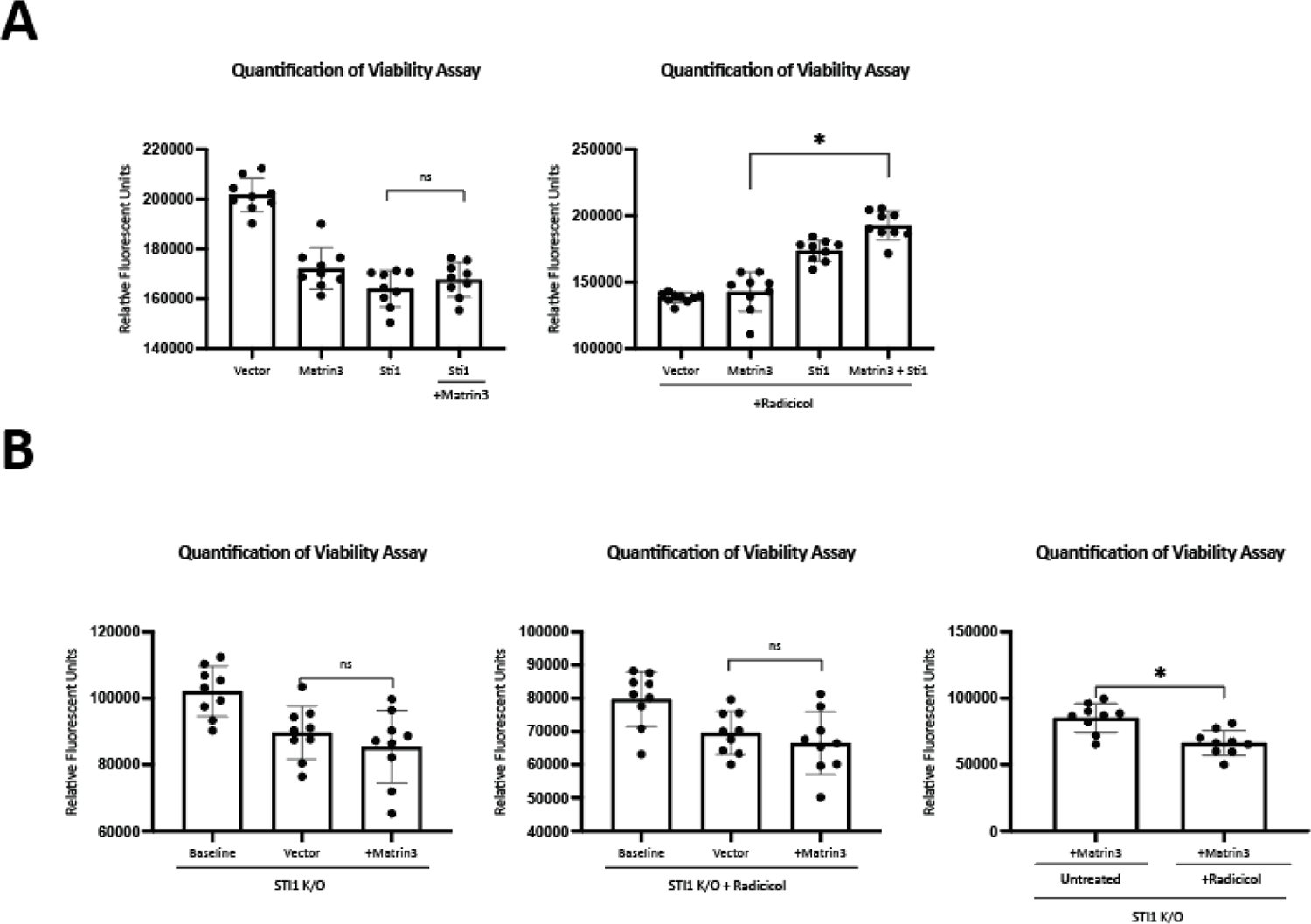
Matrin3 toxicity is altered by Sti1 and Hsp90 Inhibition in neuronal cells. **(A)** Viability assays of SN56 cells overexpressing Matrin3 with overexpressed Sti1in the absence and presence of radicicol were monitored (* = *p* < 0.05). **(B)** Viability assays of SN56 Sti1 knockout cells with overexpressed Matrin3 (* = *p* < 0.05).

## Discussion

Matrin3 has been implicated in several neurological disorders, most recently in ALS (Johnson et al., 2014, Boehringer et al., 2017, Ramesh et al., 2020, Marangi et al., 2017, Xu et al., 2016, Lin et al., 2015, Leblond et al., 2016). A total of 30 ALS associated mutations in MATR3 have been identified (Johnson et al., 2014, Boehringer et al., 2017, Senderek et al., 2009, Lin et al., 2015, Xu et al., 2016, Marangi et al., 2017, Leblond et al., 2016, Origone et al., 2015). To explore Matrin3 misfolding and toxicity, we established a yeast model expressing wild type human Matrin3 and the ALS-associated variant F115C. Similar yeast models for ALS proteins, such as TDP-43 and FUS, have delivered profound insights into mechanisms of protein misfolding and toxicity (Johnson et al., 2008, Armakola et al., 2011, Sun et al., 2011, Kryndushkin and Shewmaker, 2011, Braun et al., 2011, Jackrel et al., 2014, Liu et al., 2017, Leibiger et al., 2018, Fushimi et al., 2011, Ju et al., 2011, Daigle et al., 2013, Kryndushkin et al., 2011). Yet compared to most other ALS proteins, Matrin3 is relatively understudied and its role in neurodegeneration is poorly understood. Thus, Matrin3 presents an excellent candidate for unbiased studies in yeast that do not rely on or require substantial knowledge of protein function.

Matrin3 is similar to many other ALS proteins as it binds to RNA and localizes to NCIs. Subcellular mislocalization is a recurring pathological hallmark in ALS, which can lead to a potential loss of function of a protein in the nucleus and to toxic gain of function in the cytoplasm (Zhao et al., 2018). For other ALS associated proteins, such as TDP-43 and FUS, misfolding events are driven by their prion-like domains, where the majority of known ALS mutations are located. Intrinsically disordered domains in Matrin3 contain all known ALS-associated amino acid substitutions (Malik et al., 2018). This may suggest that the intrinsically disordered domains drive protein misfolding and ALS pathology. Our data in the Matrin3 yeast model and mammalian cells document that overexpressed Matrin3 indeed can misfold and form cytosolic and nuclear inclusions. We also found that Matrin3 misfolding and inclusion correlates with cellular toxicity in a dose-dependent manner. Further, our genetic studies revealed that Hsp90 and its co-chaperones are genetic regulators of Matrin3 toxicity and inclusion formation. Particularly, Sti1 (HOP in humans), has a strong capacity to alter Matrin3 toxicity in yeast and mammalian neuronal cell models.

Our results reveal that high expression of wild type Matrin3 and F115C is very toxic in yeast, whereas lower expression produces a milder growth defect. We further demonstrate that both RNA recognition motifs are required for toxicity in yeast, which is accordance to studies in mammalian cells (Malik et al., 2018). This indicates that Matrin3 toxicity in yeast and mammalian cells both require the RNA binding domains to cause toxicity. Further, our data show that wild type Matrin3 and F115C form inclusions in yeast even in the absence of either RRM or ZNF binding motifs. This data suggest that while RNA binding capacity of Matrin3 may underpin its toxicity it may not directly contribute to its mislocalization. This presents an example of uncoupling toxicity from inclusion formation as previously observed for TDP-43 an FUS (Guo et al., 2011, Barmada et al., 2010, Yamashita et al., 2014, Kitamura et al., 2016). Matrin3 regulates splicing events in metazoans, however, splicing mechanisms in yeast are not conserved, only very few yeast RNAs are spliced, and yeast do not express a Matrin3 homolog. Additionally, yeast do not possess the nuclear lamina proteins where Matrin3 is primarily localized in mammalian cells. This suggests that Matrin3 toxicity in yeast is based on a gain of toxic function and not a loss of function mechanism.

Neurons are tasked with constantly maintaining homeostasis under additional assaults that may directly contribute to or additively exacerbate protein misfolding (Webster et al., 2017). As shown by our data, protein misfolding stress strongly exacerbates Matrin3 toxicity in yeast and mammalian cells. Reduced Hsp90 and Hsp90 co-chaperone expression also exacerbate Matrin3 toxicity. Interestingly, inclusion formation is strongly reduced in yeast cells with reduced Hsp90 levels. Inhibition of Hsp90 in mammalian cells also result in a significant decrease in nuclear Matrin3 foci. Overexpressing Matrin3 leads to an even greater loss of inclusion formation. This suggests that Matrin3 localization is sensitive to even partial destabilization of the Hsp90 system and shows that, as shown for many misfolded proteins before, there is no strict correlation between inclusion formation and toxicity (Arrasate et al., 2004, Doi et al., 2013, Takahashi et al., 2008, Saudou et al., 1998, Miller et al., 2011, Nagai et al., 2007, Gutekunst et al., 1999, Kuemmerle et al., 1999, Skibinski and Boyd, 2012, Reiner et al., 2007, Simeoni et al., 2000, Adachi et al., 2001, Abel et al., 2001, Ordway et al., 1997, Klement et al., 1998, Romero et al., 2008, Kim et al., 1999, Taylor et al., 2003, Todd and Lim, 2013). The effect of Hsp90 inhibition on Matrin3 localization is very specific as induction of oxidative stress and proteasome inhibition do not reduce inclusion formation. Inhibition of the proteasome shows the opposite effect of Hsp90 inhibition by increasing Matrin3 inclusion formation in the nucleus. This may indicate that Hsp90 has a role in sequestering Matrin3 into inclusion. Striking the perfect balance of Hsp90 activity appears crucial as overexpressing Hsp90 also enhances Matrin3 toxicity without affecting inclusion formation. Loss or reduced expression of Hsp90 co-chaperones also exacerbates Matrin3 toxicity and reduces inclusion formation, possibly by stabilizing toxic conformations as determined before for the misfolded tau protein (Karagöz et al., 2014, Shelton et al., 2017, Weickert et al., 2020, Tortosa et al., 2009). We recapitulate these findings in mammalian cells where loss of Sti1 exacerbates Matrin3 toxicity when Hsp90 is inhibited. Moderate overexpression of Sti1 but not Aha1or Cdc37 reduces Matrin3 toxicity in yeast and neuronal cells when Hsp90 is inhibited, documenting the validity of our findings in yeast. Of note, our previous studies showed that Hsp90 and Sti1 also alter TDP-43 inclusion formation and toxicity, indicating a prominent role of these chaperones in ALS-associated protein misfolding (Lin et al., 2015).

In sum, we established and characterized a yeast model expressing human Matrin3 that allows studying Matrin3 toxicity, misfolding, mislocalization, and its interactions with cellular protein quality control systems. We show for the first time that Hsp90 and its co-chaperones have a strong capacity to modify both Matrin3 toxicity and inclusions formation. We found that Sti1 overexpression can reduce Matrin3 toxicity, indicating a specific interaction of this Hsp90 co-chaperone with Matrin3. Our data also demonstrate that there no simple positive correlation between inclusion formation and toxicity and that at least under some experimental conditions loss of inclusions correlates with increased toxicity, indicating that some inclusions may be protective. Future experiments will have to clarify the molecular mechanisms by which the Hsp90 systems and Sti1 determine Matrin3 toxicity and its role in ALS pathogenesis.

## Materials and Methods

### Materials

All chemicals and reagents were purchased from Sigma-Aldrich (Saint Louis, MI, USA) and VWR (Radnor, PA, USA). *S. cerevisiae* were grown on agar plates (20g agar/L) and in liquid media containing all essential nutrients aside from amino acids required to maintain plasmid selectivity (His-, Trp-, Leu-, or Ura-) and 2% (w/v) glucose or galactose under induced and non-induced conditions (Weinhandl et al., 2014).

All yeast strains are W303 Mat a or BY Mat α background and their derivatives (Table 1). All gene deletion strains are in the BY Mat a background obtained from gene deletion library (Thomas and Rothstein, 1989).

Plasmids for mammalian cell transfection were generated using Gateway cloning technology as previously described (Invitrogen, Carlsbad, CA, USA) (Katzen, 2007). DNA templates, pcDNA-Matrin3 and pcDNA-ΔRRM1-Matrin3, pcDNA-ΔRRM2-Matrin3, pcDNA-ΔZnF1-Matrin3 and pcDNA-ΔZnF2-Matrin3 for the generation of all constructs used in this study were generously provided by Dr. Park and Dr. Malik. (Malik et al., 2018).

Neuro2A and SNC-56 cell lines were grown in DMEM (4g/L glucose), or DMEM (1g/L glucose) (Gibco) and supplemented with 10% Bovine serum (Wisent, Saint-Bruno, Quebec, Canada), Pen/strep (Gibco), and 1% L-glutamine (Gibco). Cells were detached with warmed PBS. All wash steps were carried out using cell culture grade PBS (Gibco). Lipofectamine® 2000 (Invitrogen) was used for transfections according to the manufacturer’s instructions.

## Methods

### Yeast Transformation

Yeast strains were transformed using the LiOAc procedure as previously described (Gietz, 2014). Transformation plates were placed at 30°C for three days and single colonies were isolated and streaked (four colonies per transformation) and allowed to grow for three days. Selection by amino acid exclusion was maintained at all times. Plates were not allowed to age more than three weeks before re-streaking to maintain fresh strains.

### Spotting Assays

Spotting assays were performed as previously described (Di Gregorio et al 2020). In brief, yeast strains were inoculated overnight in 3ml of selective growth media at 30°C and grown to saturation. The optical density (OD600) was measured for each strain following 12h-16h of growth. Cultures were transferred to the top wells of sterile 96-well plates (Greiner bio-one #650161) and normalized to a starting OD600 of 0.1 in 200ul. A five-fold serial dilution was performed and cells plated. In the experiments using chemical treatments, cells were plated on agar plates containing the indicated concentration of various chemical stress inducers (1mM AZC, 50µM MG132, 100uM H_2_0_2_, 25µM radicicol, 2.5µg/ml tunicamycin, 5mM DTT).

### Quantification

Plates with yeast spotting assays were photographed on Bio-Rad GelDoc system (white light setting). Images were first processed in Photoshop and converted into black and white images. Images were then imported into ImageJ for quantification. GraphPad prism was used to generate bar graphs. A detailed protocol for the quantification of yeast growth assays is provided under https://star-protocols.cell.com/protocols/166

### Microscopy

Following a minimum of 16 hrs induction, yeast cells were pipetted onto glass slides and sealed under coverslips. Fluorescence microscopy was performed using the Cytation5 cell imaging multi-mode reader (BioTek, Winooski, VT, USA). The following LED cubes and imaging filter cubes from biotech were employed for GFP tagged proteins: 465 LED 1225001 Rev J, GFP 469/525 1225101 Rev J, BioTek; and for YFP tagged proteins: 465 LED 1225001 Rev J, CFP 445/510 1225107 Rev I. Images were captured on the 20x objective. Image analysis was completed using the Gen5 imaging software (BioTek).

Yeast microscopy data was quantified as previously described (Di Gregorio et al., 2020).

### Mammalian cell culture

#### Transfection

We used mouse wild type SN56 and Sti1 knock out (Lackie et al., 2020)), and mouse derived (Neuro2a) lines. Partial differentiation of Neuro2a and SN56 cells was carried out by growing cells in serum reduced media. Transfection of Matrin3 WT, ΔRRM1, ΔRRM2, ΔZnF1, and ΔZnF2 and DNAJ A1, STi1 SiRNA were carried out with Lipofectamine® 2000 according to the manufactures protocol.

#### Viability assays

Neuro-2a and SNC-56 cells were seeded at 3,000 cells/well in 96 white well plates and transfected as stated above. The Cell Titer Glo 2.0 (Promega), and the luciferase based luminescence viability assay was used to detect viable cells. Luminescence was measured on Cytation5 plate reader (BioTek).

#### Immunofluorescence microscopy

Prior to imaging, cells were seeded at 800-1,000 cells/well in 6 well dishes on coverslips in DMEM, 5% or 0% bovine serum. Cells were probed with anti-Matrin3 (Sigma HPA036565), or anti-Hsp90 (ab13492) overnight. Alexa Fluor 488 or Alexa Fluor 688 secondary antibodies (Thermo-Fischer Scientific) were utilized for fluorescent microscopy. Images were taken on the Cytation5 as described for yeast. Confocal immunofluorescence microscopy was performed on the Zeiss LSM800 with Airyscan

### Western blot and SDD-AGE

#### Yeast protein lysis

Yeast cells were inoculated in 3ml of non-inducing media and allowed to grow overnight at 30°C. The following evening, cultures were spun and washed twice in water before resuspension in 10ml inducing growth media and returned to 30°C for overnight growth. Following 16 hours of induction, cells were lysed using the alkaline lysis method as previously described (Kushnirov, 2000).

SDS Page and Western Blot: Protein samples in loading buffer (2x TAE, 20% glycerol, 8% SDS, bromophenol blue) were loaded on a Criterion™ precast 4%–12% Tris-HCl gel (Bio-Rad) and run at 110 V. The proteins were semi-dry-transferred to a membrane for 7 min at 35 V. The membrane was blocked in 2% BSA in PBS-Tween (0.01%) for a minimum of 1 h. The blot was incubated with primary antibody in the same solution overnight at 4°C. The membrane was washed five times with PBS-Tween (0.01%) before secondary antibody was added (anti-mouse Alexa 680), in 2% BSA in PBS-Tween (0.01%). The membrane was incubated at room temperature for 1 h before it was washed five times again with PBS-Tween (0.01%). Antibodies were visualized with the ChemiDoc MP imaging system (Bio-Rad) and analyzed using the Image Lab 5.2 software (Bio-Rad.) Anitbodies: Anti GFP (Sigma), Anti FLAG (Sigma F3040)

SDDAGE: Samples were normalized using BCA for total protein concentrations and diluted in loading buffer (2x TAE, 20% glycerol, 8% SDS, bromophenol blue). Samples were loaded into 1% SDS, 1.8% agarose gel and run in 1% SDS, 1xTAE buffer for two hours at 80 volts. Protein transfer was conducted using the TurboBlotter and blotting stack apparatus (Whatman) by gravity and capillary force using TBS transfer buffer (10mM Tris pH [7.5], 154mM NaCl), on nitrocellulose membrane. Protein was detected by standard western blotting (see above).

## Statistics

Biologicals in triplicate are quantified as stated above. Mammalian cell line viability assay represent an n=9 for each condition. Statistical analysis was carried out applying One Way Analysis of Variance, Tukey Post Hok using IBM SPSS. Error bars represent standard deviation.

## Acknowledgments

We would like to thank Dr. Park (University of Toronto) & Dr. Baramada (University of Michigan) for providing the Matrin3 vectors.

## Author Contributions statement

All experiments were conducted by S.E.D., M.A.E., A.R. and A.S.. Experimental design was conducted by SED and MLD. Analyses of data was conducted by MLD, SEG and data interpretations by SED, and MLD. Manuscript preparation, editing and proofreading was conducted by SED R.J.R, and MLD.

## Supplementary Figures

**Supplementary Figure 1.**
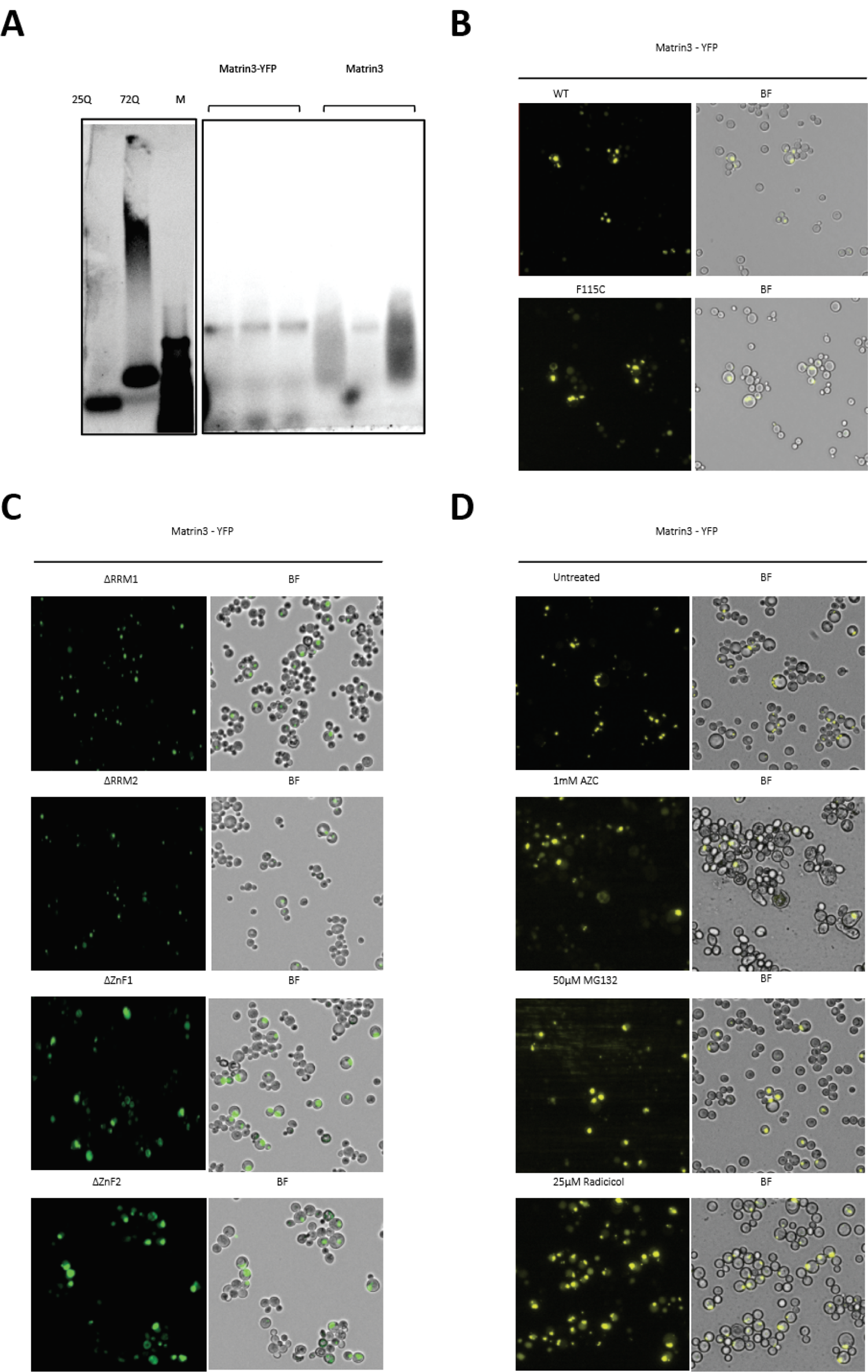
**(A)** Semi-denaturing detergent gel electrophoresis (SDD-AGE) of protein lysates of yeast cells expressing Matrin3 or polyglutamine expansion proteins. **(B)** Expanded fields of fluorescent microscopy of Matrin3-YFP and F115C-YFP expressing yeast cells corresponding to Figure 1C. **(C)** Expanded fields of fluorescent microscopy of Matrin3-YFP ΔRRm1, ΔRRM2, ΔZnF1, and ΔZnF2 truncations expressing yeast cells corresponding to Figure 1E. **(D)** Expanded fields of fluorescent microscopy of Matrin-YFP expressed in untreated yeast cells and cells treated with 1mM AZC, 50µM MG132, 25µM Radicicol corresponding to Figure 2H.

**Supplementary Figure 2.**
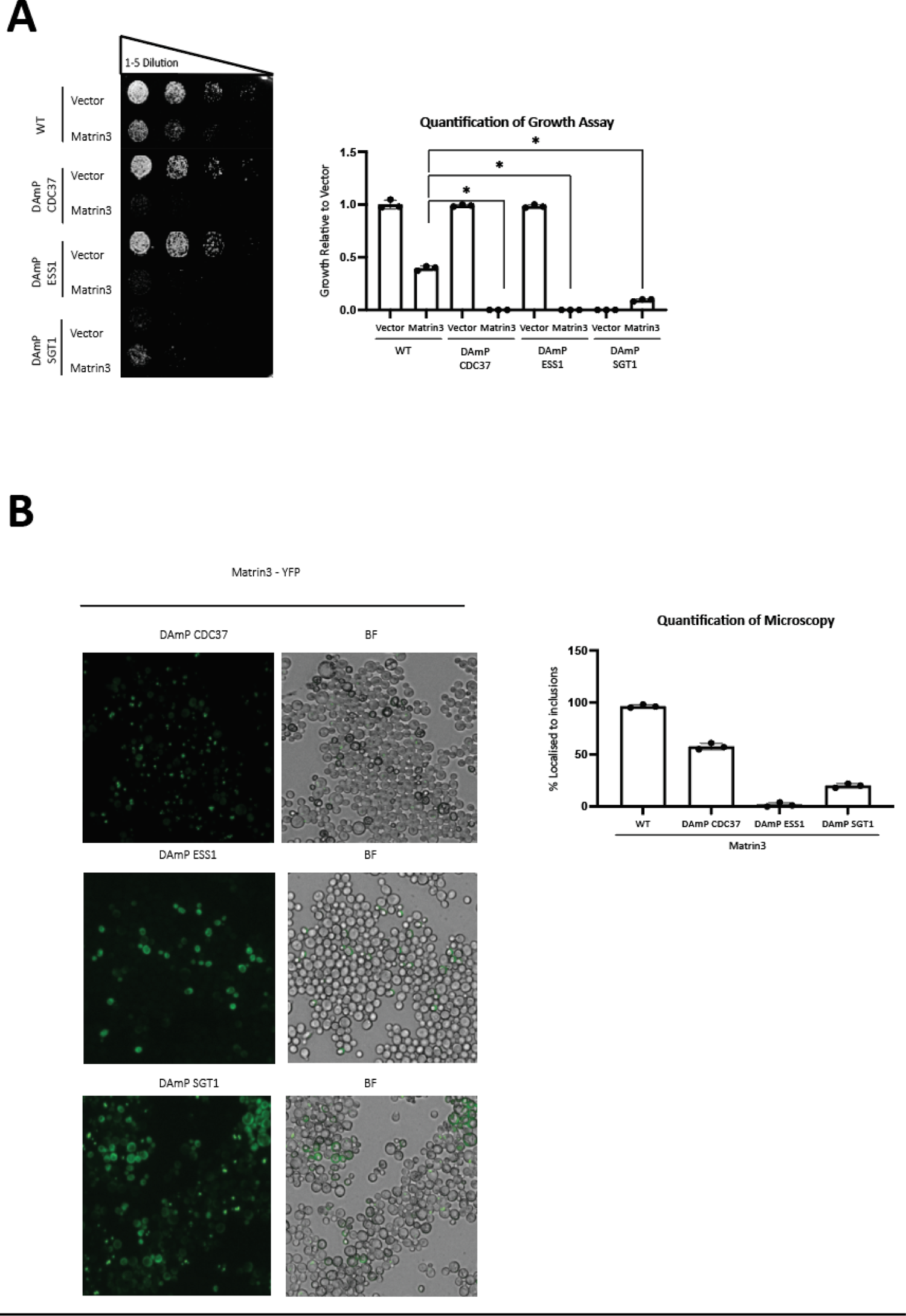
**(A)** Growth assay of yeast cell expressing Matrin3 (low expression system) in wild type and DAmP yeast strains of essential Hsp90 co-chaperones as indicated (* = *p* < 0.05). **(B)** Expanded fluorescent microscopy fields yeast cells expressing of Matrin3-YFP in the indicated DAmP yeast strains of essential Hsp90 co-chaperones and quantification of Matrin3-YFP inclusions (* = *p* < 0.05).

**Supplementary Figure 3.**
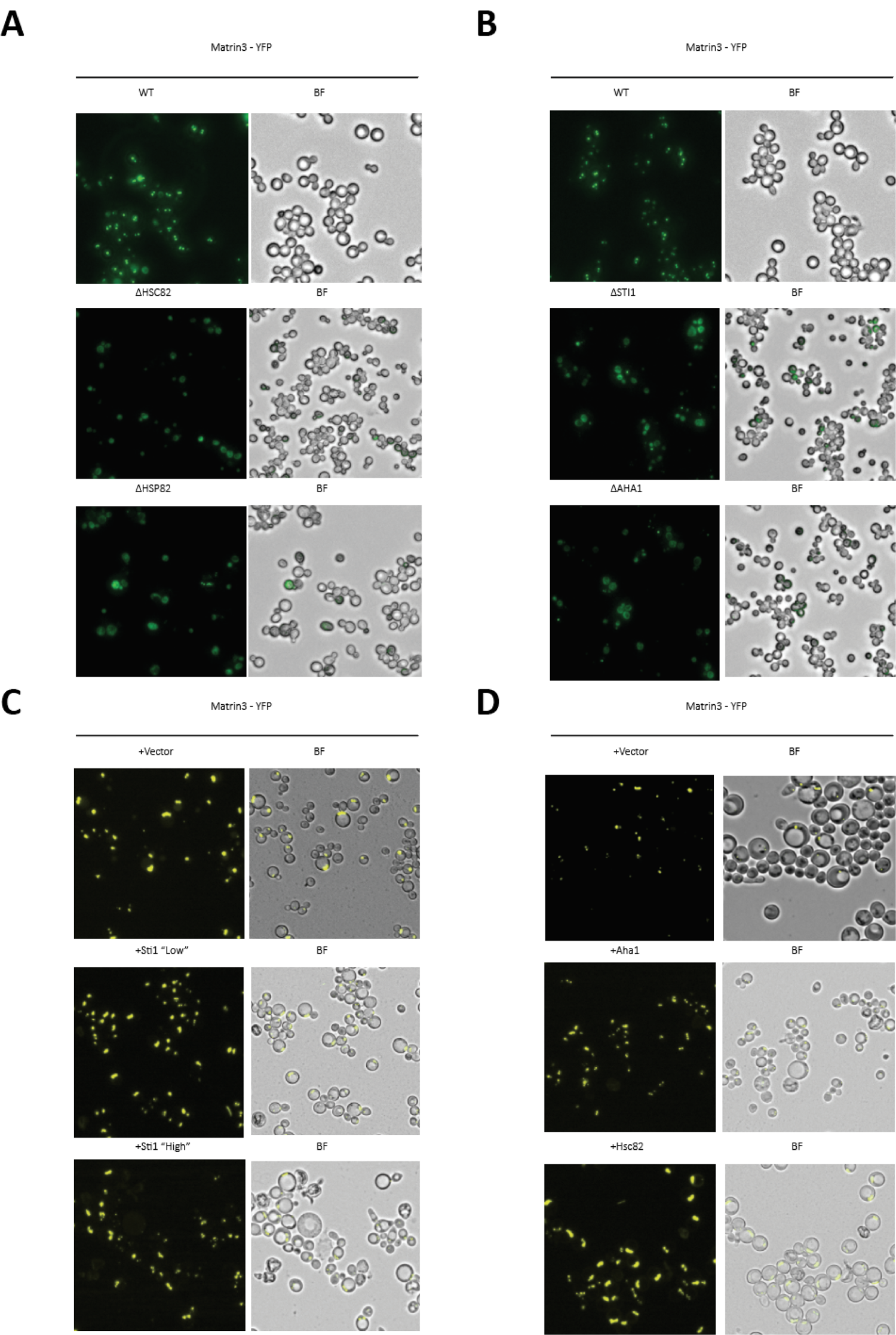
**(A)** Expanded fluorescent microscopy fields of Matrin3-YFP WT expressed in wild type yeast strains deleted for HSC82 (Δhsc82) and HSP82 (Δhsp82) corresponding to Figure 4C. **(B)** Expanded fluorescent microscopy fields of Matrin3-YFP expressed in the wild type and yeast deletion strains, Δsti1 and Δaha1 corresponding to Figure 5C. **(C)** Expanded fluorescent microscopy fields of Matrin3-YFP expressed in yeast cells overexpressing Sti1 (‘low’ overexpression), and (‘High’ overexpression) and vector control corresponding to Figure 6A and 6B. **(D)** Expanded fluorescent microscopy fields of Matrin3-YFP expressed in yeast cells overexpressing Aha1 and Hsc82 and vector control corresponding to Figure 6C.

**Supplementary Figure 4.**
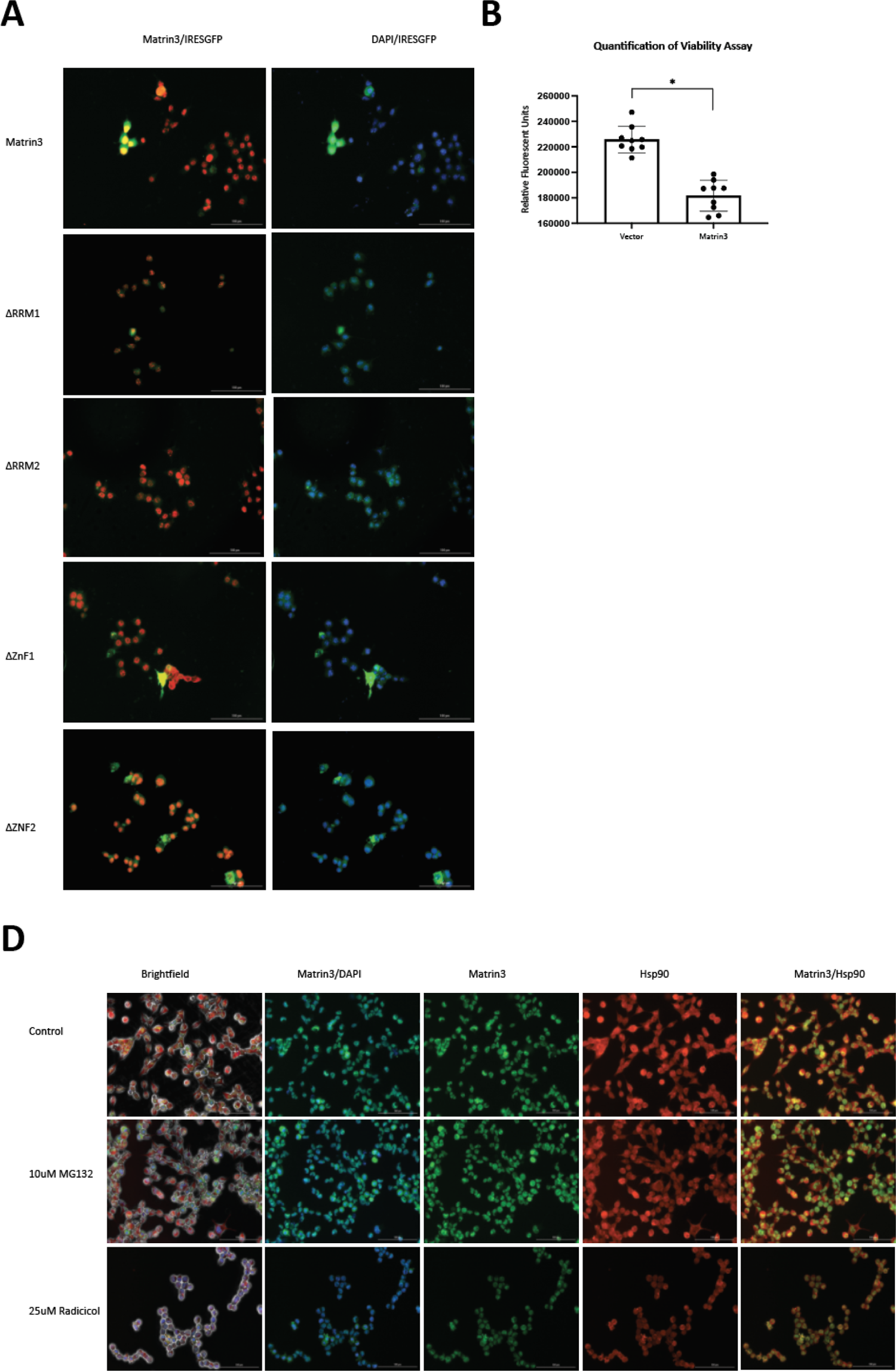
**(A)** Immunofluorescence microscopy of N2a cells transfected with expression vectors for Matrin3 WT and ΔRRM1, ΔRRM2, ΔZnF1, and ΔZnF2 truncations. **(B)** Viability assays of N2a cells transiently transfected with Matrin3 expression vectors (* = *p* < 0.05). (C Immunofluorescence microscopy of N2a cells detecting endogenous Matrin3 and Hsp90 in untreated and cells treated with MG132 or radicicol.

**Supplementary Figure 5.**
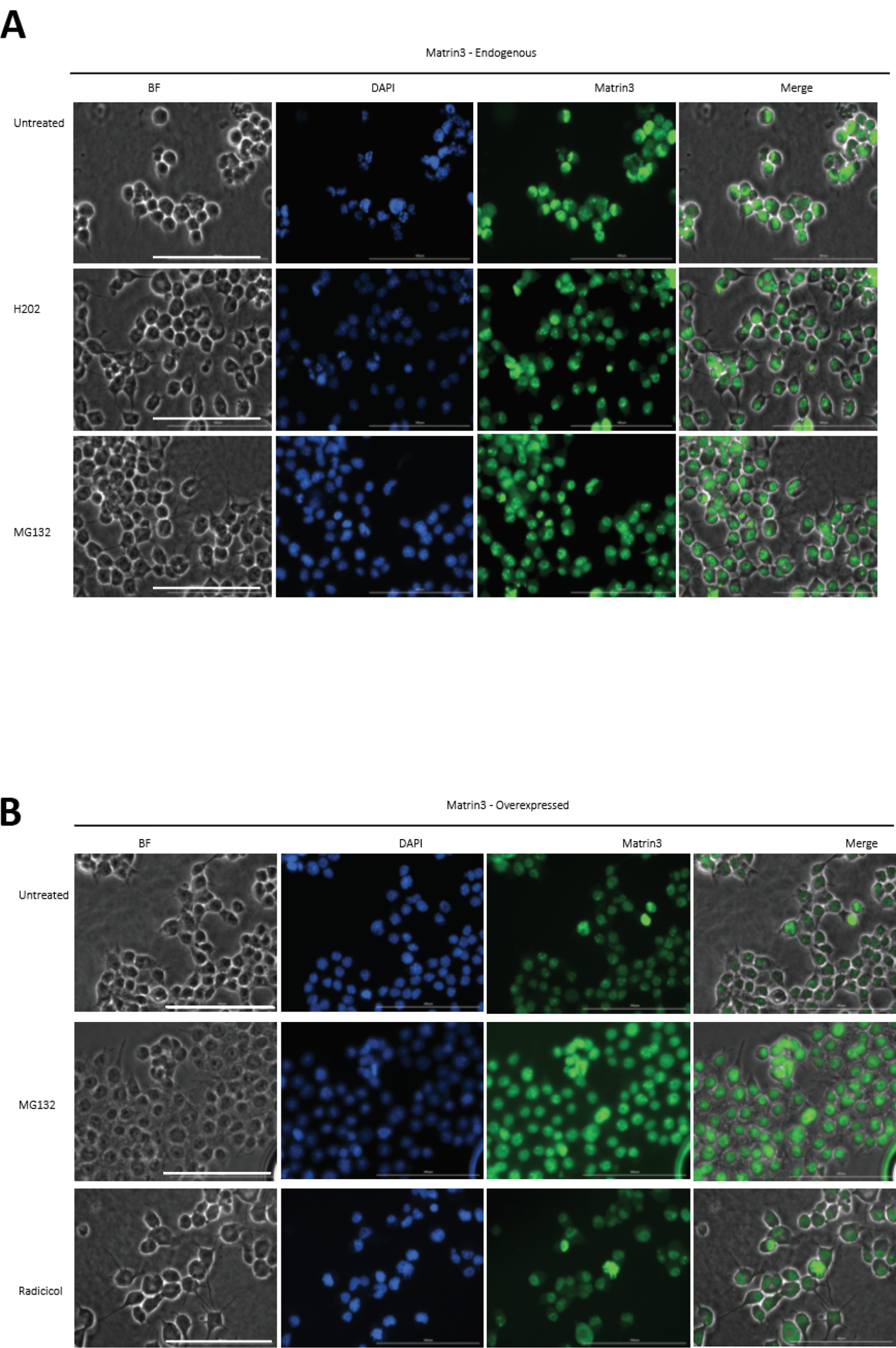
**(A)** Immunofluorescence microscopy of N2a cells detecting endogenous Matrin3 treated with H_2_0_2_, MG132, or radicicol, and untreated controls. **(B)** Immunofluorescence microscopy of N2a cells transiently transfected with Matrin3 expression vectors treated with H_2_0_2_, MG132, or radicicol or untreated controls.

**Supplementary Figure 6.**
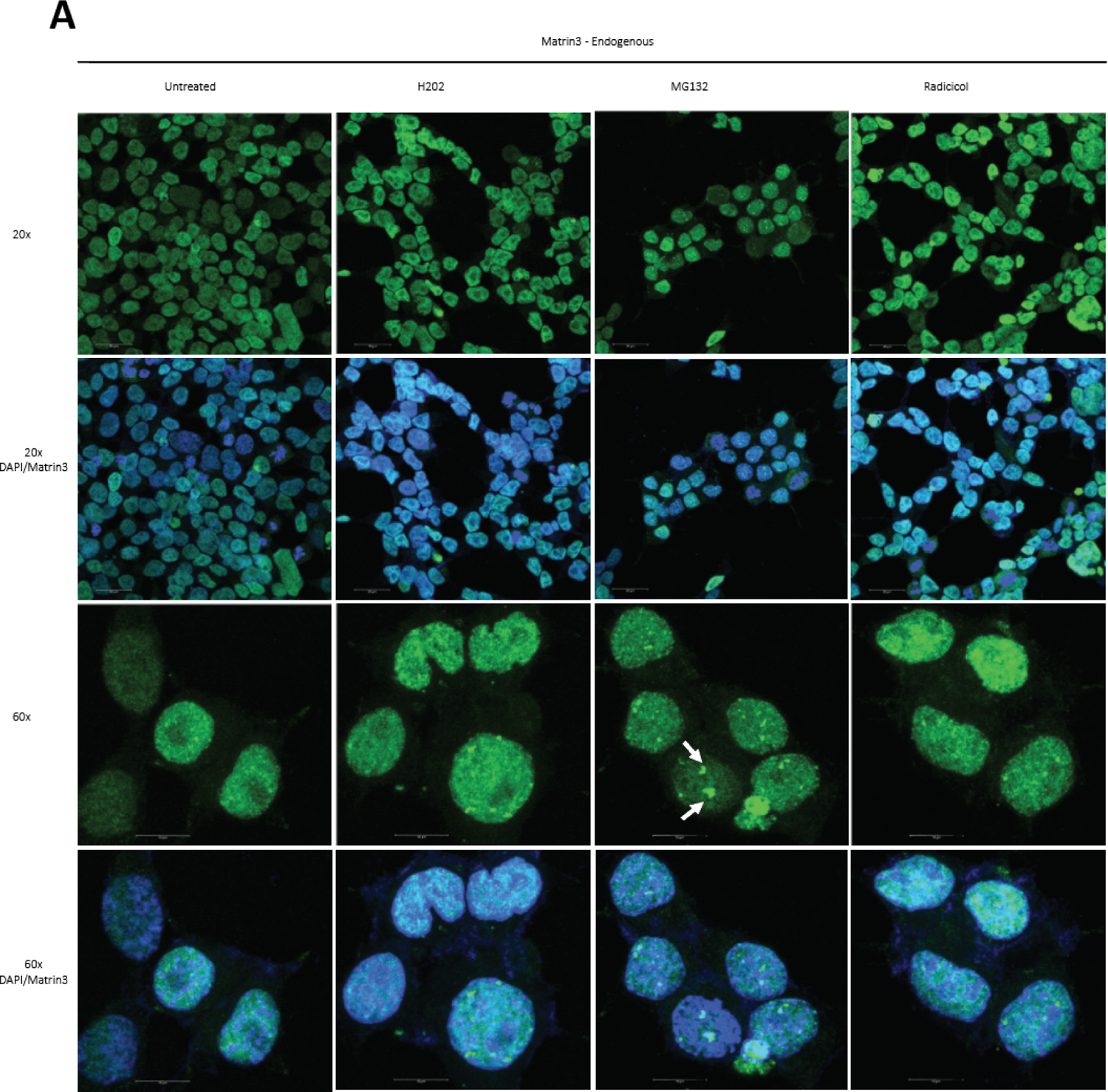
**(A)** Immunofluorescence microscopy of N2a cells detecting endogenous Matrin3 treated with H_2_0_2_, MG132, or radicicol, and untreated controls on 20x and 60x objectives.

**Supplementary Figure 7.**
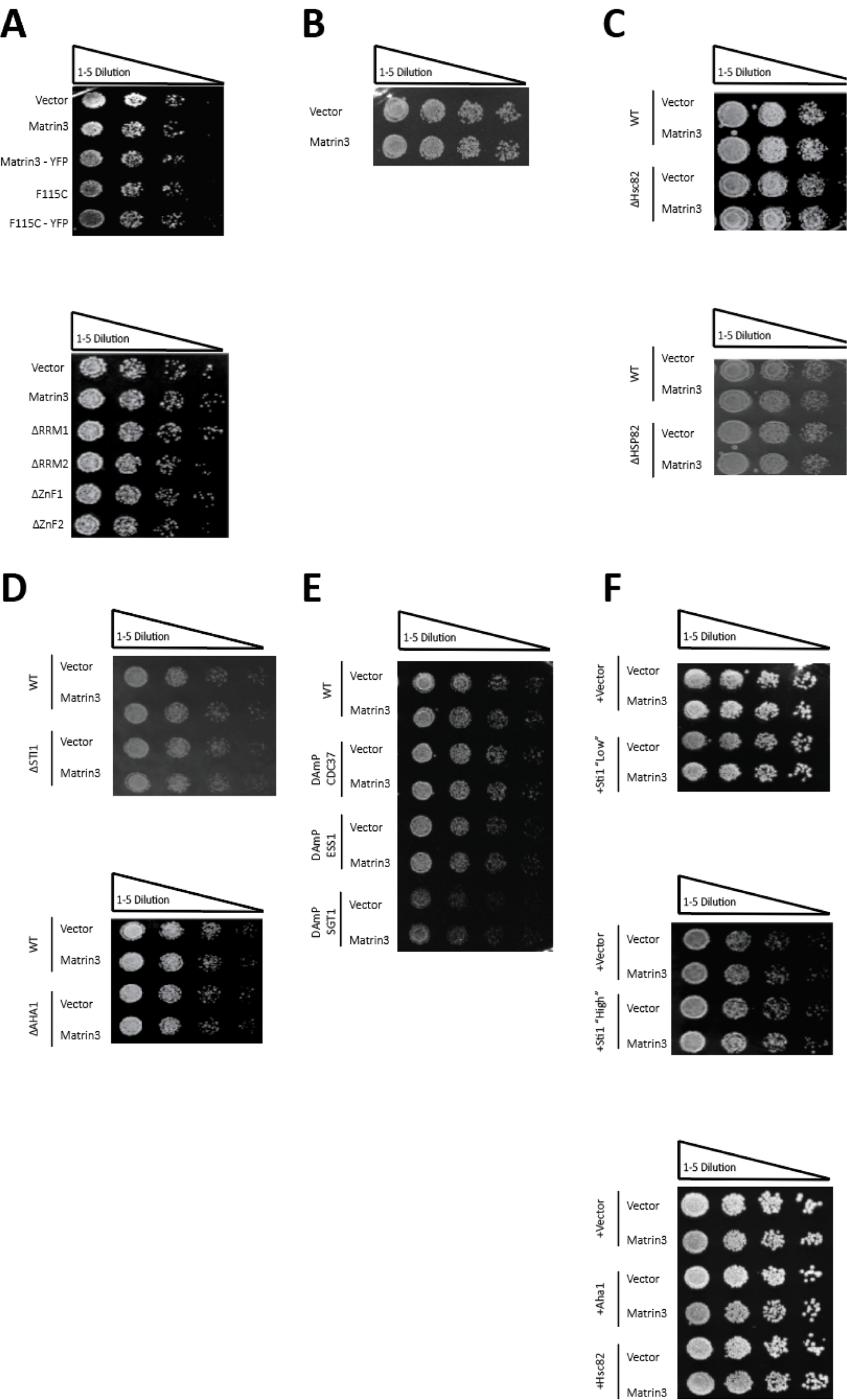
**(A)** Non-inducing control plates corresponding to Figure 1. **(B)** Non-inducing control plates corresponding to Figure 2. **(C)** Non-inducing control plates corresponding to Figure 3. **(D)** Non-inducing control plates corresponding to Figure 4. **(E)** Non-inducing control plates corresponding to Supplementary figure 2. **(F)** Non-inducing control plates corresponding to Figure 5.

**Supplementary Figure 8.**
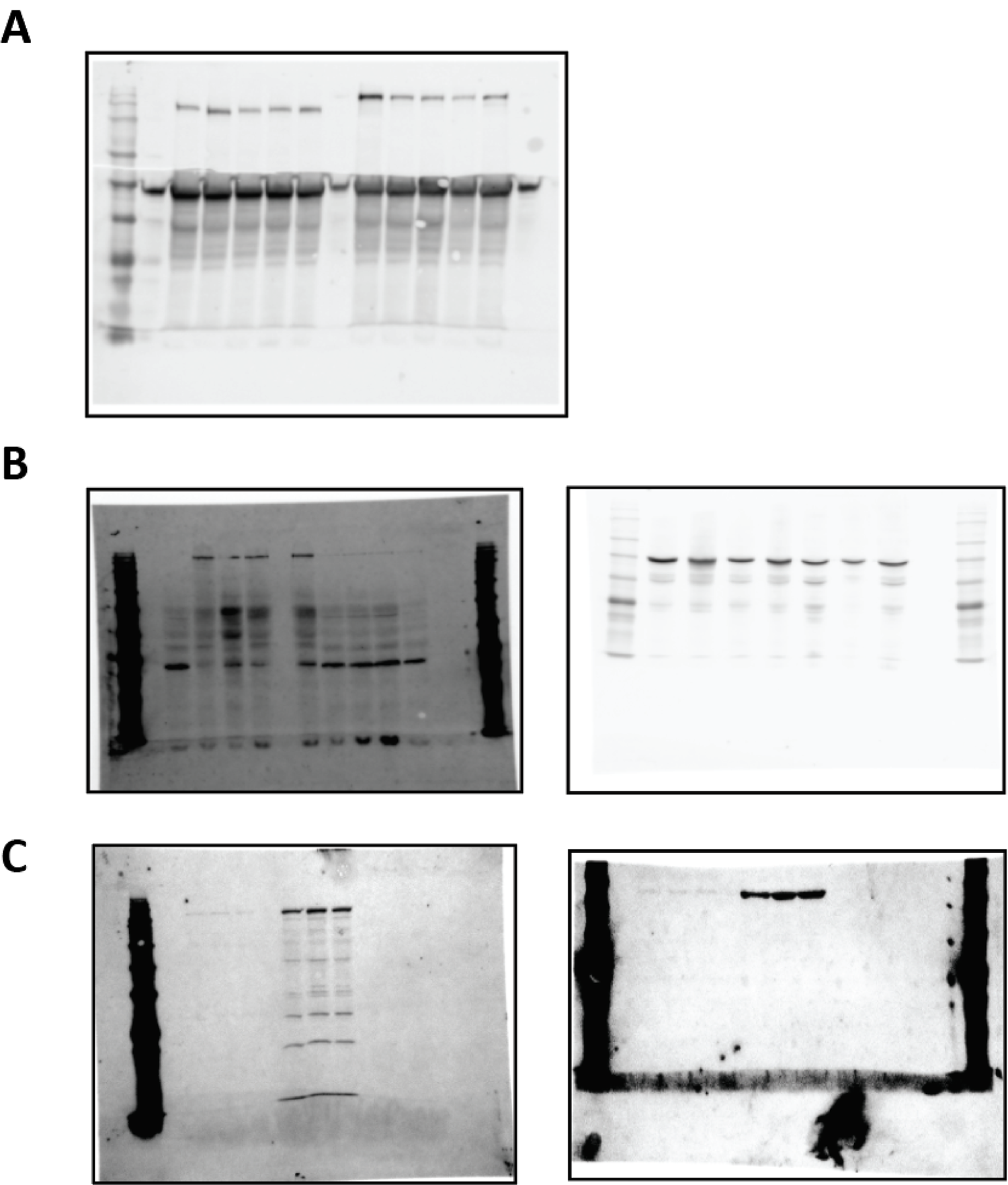
**(A)** Uncropped western blot corresponding to Figure 1F. Untagged set (left), and matching YFP-tagged set (right). **(B)** Uncropped western blots pertaining to Figure 2I. Anti-Matrin3 blot (left), anti-PGK1 blot (right). **(C)** Uncropped western blots pertaining to Figure 2I. Anti-Matrin3 blot (left), anti-Tubulin blot (right).

